# Disruption of the autism-associated gene *SCN2A* alters synaptic development and neuronal signaling in patient iPSC-glutamatergic neurons

**DOI:** 10.1101/2021.09.14.460368

**Authors:** Chad O. Brown, Jarryll Uy, Nadeem Murtaza, Elyse Rosa, Alexandria Alfonso, Sansi Xing, Biren M. Dave, Savannah Kilpatrick, Annie A. Cheng, Sean H. White, Jennifer Howe, Stephen W. Scherer, Yu Lu, Karun K. Singh

## Abstract

*SCN2A* is an autism spectrum disorder (ASD) risk gene and encodes a voltage-gated sodium channel. However, the impact of autism-associated SCN2A *de novo* variants on human neuron development is unknown. We studied SCN2A using isogenic *SCN2A*^-/-^ induced pluripotent stem cells (iPSCs), and patient-derived iPSCs harboring a p.R607* or a C-terminal p.G1744* *de novo* truncating variant. We used Neurogenin2 to generate excitatory glutamatergic neurons and found that *SCN2A*^+/*p*.*R607*\*^ and *SCN2A*^-/-^ neurons displayed a reduction in synapse formation and excitatory synaptic activity using multielectrode arrays and electrophysiology. However, the p.G1744* variant, which leads to early-onset seizures in addition to ASD, altered action-potential dynamics but not synaptic activity. Proteomic and functional analysis of *SCN2A*^+/*p*.*R607*\*^ neurons revealed defects in neuronal morphology and bioenergetic pathways, which were not present in *SCN2A*^+/*p*.*G1744*\*^ neurons. Our study reveals that SCN2A *de novo* variants can have differential impact on human neuron function and signaling.

**HIGHTLIGHTS:**

- Isogenic *SCN2A*^-/-^ neurons display intrinsic hyperexcitability and impaired excitatory synapse function
- *SCN2A*^+/*p*.*R607*\*^ variant reduces excitatory synapse function in patient neurons
- C-terminal *SCN2A*^+/*p*.*G1744*\*^ variant enhances action potential properties but not synaptic transmission in patient neurons
- *SCN2A*^+/*p*.*R607*\*^ variant display impacts on morphological and bioenergetic signaling networks through proteomic and functional analysis

**eTOC:**

- Brown et al. examined Autism-associated *SCN2A* variants using patient-derived iPSC NGN2-neurons. They discover that genetic variants differentially impact neuronal development and synaptic function, and highlight neuronal and bioenergetic signaling networks underlying SCN2A loss-of-function.

## INTRODUCTION

Autism spectrum disorder (ASD) is a childhood-onset, heterogeneous group of neurodevelopmental disorders that can have a range of severities between individuals. ASD is characterized by deficits in social communication, restricted and repetitive patterns of behavior, or interests (Ofner et al., 2018). Large-scale genetic sequencing studies have highlighted the contributions of genetic variants in hundreds of genes to the underlying etiology, including rare-inherited and *de novo* variants being a major contributor of risk (Bai et al., 2019; Grove et al., 2019; Iossifov et al., 2014; Ruzzo et al., 2019; Sanders et al., 2015; Satterstrom et al., 2020; Yuen et al., 2017). Whole exome and genome sequencing studies of ASD cohorts have identified multiple *de novo* genetic variants within *SCN2A*, which has led to *SCN2A* being one of the strongest individual candidate risk genes for ASD (Crawford et al., 2021; Sanders et al., 2012; Satterstrom et al., 2019, 2020; Wang et al., 2016). *SCN2A* encodes the neuronal α-subunit of the voltage-gated sodium channel Na_V_1.2. which is predominately expressed in excitatory glutamatergic neurons within the cortex. The function of SCN2A and its location changes throughout development, where early expression at the distal axon initial segment (AIS) is essential for initiation and propagation of action potentials; while later in development, expression is clustered at the proximal AIS, where *SCN2A* function regulates action potential backpropagation (Bender and Trussell, 2012; Hu et al., 2009; Kole and Stuart, 2012; Spratt et al., 2019). *SCN2A* is thought to predominantly regulate excitatory neuron function, while inhibitory neuron characteristics remain largely unchanged (Ogiwara et al., 2018; Spratt et al., 2019; Wang et al., 2021).

Understanding how variants in *SCN2A* contribute to neurological disorders is beneficial as it may inform potential therapies or long-term clinical outcomes. Cell line and computational studies have been instrumental in understanding the potential impact of the *SCN2A* variants (Begemann et al., 2019; Ben-Shalom et al., 2017; Echevarria-Cooper et al., 2021; Que et al., 2021). These studies have indicated that *SCN2A* gain-of-function variants are likely associated with an enhancement of channel function and epileptic encephalopathy. Loss-of-function variants in *SCN2A* including those that result in protein truncations, are associated with reduced channel function, leading to ASD and intellectual disability (Begemann et al., 2019; Ben-Shalom et al., 2017; Sanders et al., 2018). Given the association between loss-of-function *SCN2A* variants and ASD, the majority of studies have used *Scn2a*^+/-^ mice as an animal model. These mice have defects in spatial memory and learning, and social behavior (Ogiwara et al., 2018; Spratt et al., 2019; Tatsukawa et al., 2019). Additionally, Na_v_1.2 channels in deep layer (5/6) prefrontal cortical excitatory neurons mediate backpropagation of action potentials to dendrites, postsynaptic calcium influx, synaptic function, and plasticity (Spratt et al., 2021, 2019). These studies suggest heterozygous loss of Scn2a function, which models protein-truncating variants, results in developmental excitatory circuit abnormalities in the cortex. While mouse models provide important insights, it does not possess the human and/or patient’s genetic background, which can influence disease phenotypes (Mis et al., 2019). Further, patient-derived iPSC-neurons modeling missense SCN8A-associated epilepsy encephalopathy variants revealed defects in sodium channel currents and action potential dynamics, which likely drive patient phenotypes (Tidball et al., 2020). These studies indicate that the impact of sodium channel variants on human neuronal function can be revealed using an iPSC system. However, *SCN2A* has not been modeled using a patient-specific neuronal system allowing for direct comparisons with familial controls (Deneault et al., 2018; Kohlnhofer et al., 2021; Lu et al., 2019). Our research group and others previously showed that *SCN2A*^*-/-*^ glutamatergic neurons display reduced synaptic activity. However, computational, cell line, mouse and iPSC-derived neuron models have yet to show whether they reflect phenotypes from SCN2A patient-derived neurons, and its critical signaling mechanisms (Begemann et al., 2019; Ben-Shalom et al., 2017; Deneault et al., 2018; Ogiwara et al., 2018; Shin et al., 2019; Spratt et al., 2021, 2019; Tatsukawa et al., 2019; Wolff et al., 2017; Zhang et al., 2021). SCN2A is a large channel with multiple domains, and the working hypothesis is that *de novo* protein-truncating variants result in a loss of channel activity. However, given the size of SCN2A, it remains unknown whether protein-truncating variants at different locations in SCN2A have the same functional effect.

To address the gap in disease modeling of *SCN2A* variants, we generated iPSCs from two ASD probands with sex-matched parental controls, as well as a new isogenic *SCN2A*^*-/-*^ iPSC line. We used the Neurogenin2 (NGN2) expression protocol (Zhang et al., 2013) to generate glutamatergic neurons (iNeurons) for electrophysiological and proteomic studies (Figure 1A). iNeurons were generated from an ASD patient with the *SCN2A*^*+/p*.*R607\**^ truncating variant, and from an ASD individual that show comorbidity for early-onset seizures with the *SCN2A*^*+/p*.*G1744\**^ truncating variant. These variants are located near opposite ends of the channel, providing a comparison of different protein domains. We found that isogenic *SCN2A*^*-/-*^ iNeurons produce paradoxical hyperexcitable phenotypes in iNeurons, similar to Scn2a KO neurons (Spratt et al., 2021, 2019; Zhang et al., 2021). Further electrophysiological analysis of isogenic *SCN2A*^*-/-*^ and *SCN2A*^*+/p*.*R607\**^ iNeurons revealed they have a similar reduction in synapse development, synaptic transmission, and neuronal network activity. However, *SCN2A*^*+/p*.*G1744\**^ iNeurons displayed no changes in synaptic transmission, but possessed enhanced action potential characteristics. Further, *SCN2A*^*+/p*.*G1744\**^ neuronal network activity were able to compensate and recover most firing metrics in late development. To probe the downstream molecular effect of a loss of *SCN2A* function, we performed shotgun proteomics and determined that neurodevelopmental and bioenergetic pathways were altered. Further, this was functionally validated in patient neurons using metabolic assays. These findings suggest that truncating variants in *SCN2A* can result in abnormal neuronal function via differing mechanisms, highlighting the need to study multiple variants in the same ASD risk genes to understand neurodevelopmental disorder mechanisms.

**Figure 1.**
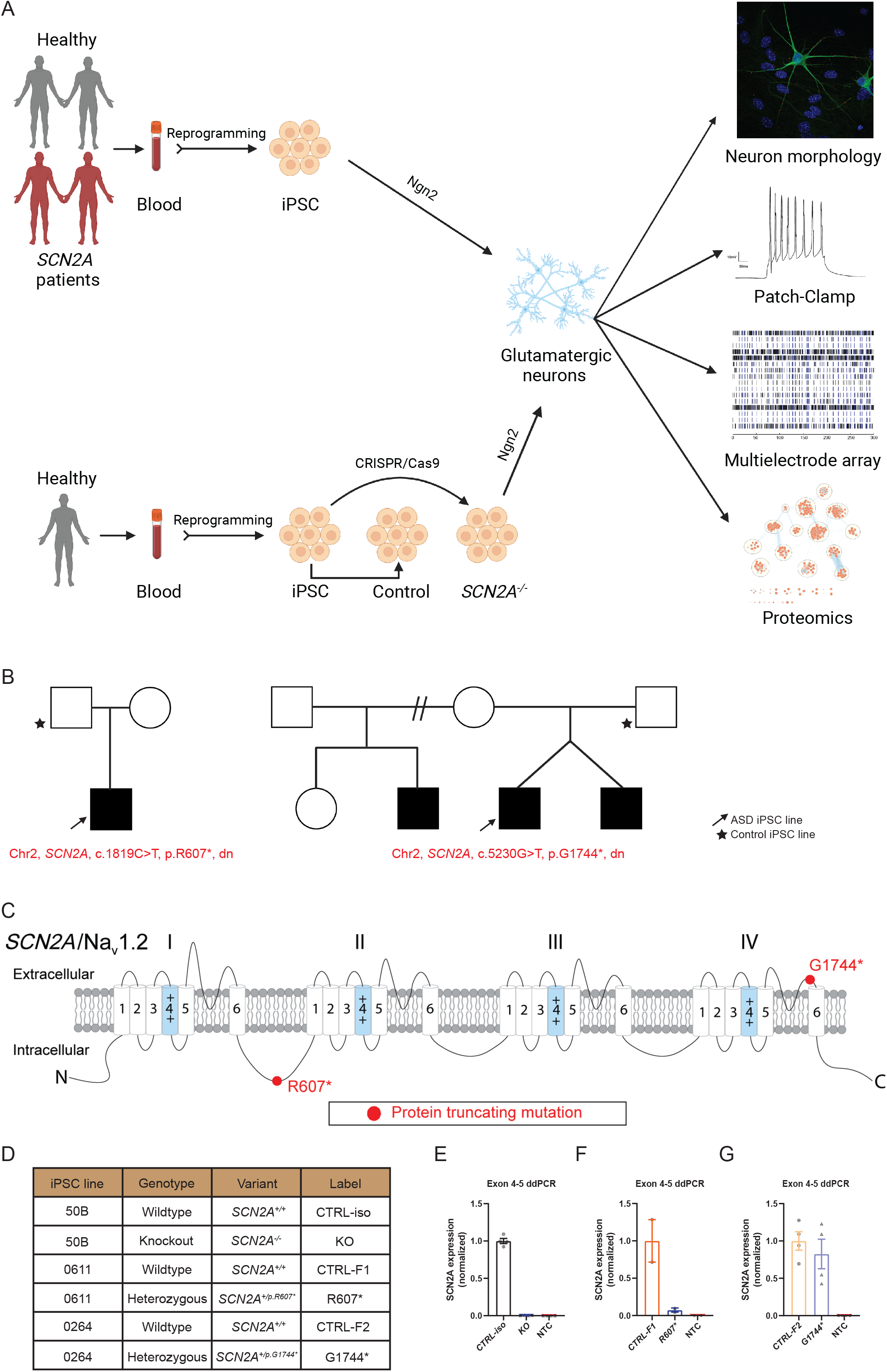
Experimental pipeline to probe cellular and molecular consequences of SCN2A deficiency in iPSC-derived iNeurons. See also figure S1. A) Experimental pipeline to test cellular and signaling function. B) SCN2A patient and family pedigrees. C) Schematic of SCN2A structure with the associated location of the *de novo* variants. D) Classification of iPSC lines used and their associated naming convention. E-G) Digital droplet PCR normalized *SCN2A* expression probed at exon 4 – 5. (E) CTRL-iso (n=4) and KO (n=4) iNeurons. (F) CTRL-F1 (n=2) and R607* (n=2) iNeurons. (G) CTRL-F2 (n=4) and G1744* (n=4) iNeurons.

## RESULTS

### Generation and characterization of isogenic SCN2A knockout and two patient-derived truncating *de novo* variants in human iPSCs and iNeurons

We recruited two unrelated families with *de novo* protein-truncating variants in *SCN2A* in the proband and sex-matched parental controls that did not have the variant (Figure 1B). The variants were identified by whole-genome sequencing from the MSSNG database (Yuen et al., 2017). Both probands were male; one had ASD and a *de novo* truncating variant at amino acid position 607 (*SCN2A*^*+/p*.*R607\**^), while the other proband had ASD and early-onset seizures with a C-terminal truncating variant at position 1744 (*SCN2A*^*+/p*.*G1744\**^) (Figure 1B). iPSCs were generated as previously described (Deneault et al., 2018, 2019), and all iPSC lines had a normal karyotype, expressed the pluripotency markers OCT4 and NANOG, and were mycoplasma free (Figure S1A and C). We confirmed that the iPSCs from the probands carried the *SCN2A* variant (Figure S1D). In addition to the patient-derived and parental control iPSCs, we generated a new *SCN2A*^*-/-*^ iPSC line using CRISPR/Cas9 to insert a STOP-tag into exon 5 of an iPSC line we previously used (named 50B) (Deneault et al., 2018), which was confirmed by sequencing (Figure S1D) (referred to as CTRL-iso and KO respective to wild-type and knockout neurons). The first family, or *SCN2A*^*+/p*.*R607\**^ variant, and familial control are referred to as CTRL-F1 and R607*, while the second family or *SCN2A*^*+/p*.*G1744\**^ variant and familial control are referred to as CTRL-F2 and G1744* (Figure 1D).

### Altered SCN2A expression by CRISPR knockout and two patient *de novo* truncating variants in human iPSC-iNeurons differentially effects neuronal activity and maturation

We used a modified Neurogenin2 induction and plating protocol (Zhang et al., 2013) to generate iNeurons. We validated *SCN2A* expression in iNeurons using digital droplet PCR (ddPCR) due to the difficulty in performing reliable and consistent western blots for SCN2A. SCN2A is a very large membrane protein and is difficult to extract in sufficient quantities. Compared to controls, we found that probes targeting exon 4-5 had minimal detectable *SCN2A* transcript in isogenic KO neurons, demonstrating that the CRISPR-Cas9-inserted 3x stop-tag single-stranded oligodeoxynucleotides donor (ssODN) disrupted SCN2A expression (Figure 1E). Using patient R607* iNeurons, the ddPCR detected a small quantity of *SCN2A* mRNA, which is likely derived from the remaining allele (Figure 1F). We also examined iNeurons from the G1744* variant located in exon 27, which is beyond the predicted non-sense mediated decay breakpoint (Nagy and Maquat, 1998; Sanders et al., 2018). It is hypothesized that *SCN2A* variants associated with ASD are loss-of-function and reduce channel activity; however, this variant could be complex given this individual has early-onset seizures, and the variant is located at the C-terminus before the calmodulin binding domain. We quantified *SCN2A* expression in G1744* iNeurons at exon 4-5 and found that the variant produces a small reduction in expression of *SCN2A* (Figure 1G). This suggests that the G1744* variant has a lesser impact on *SCN2A* mRNA levels.

Next, we examined dendrite growth and synapse formation since previous studies have shown that *Scn2a*^*+/-*^ animal studies have immature spine development in cortical excitatory neurons (Spratt et al., 2019). We used DIV 26-28 iNeurons and identified synapses by staining for the presynaptic terminal marker Synapsin1, together with MAP2, a microtubule-associated protein to outline dendrites. We did not observe statistical differences for KO, R607* or G1744* iNeurons compared to their respective controls (Figure 2A-C). However, the isogenic KO and both patient R607* and G1744* iNeurons displayed decreased Synapsin1-positive synaptic density, with fewer presynaptic terminals being formed (Figure 2D-F). The size of presynaptic terminals was increased in KO and G1744* iNeurons, but there was no change in patient R607* iNeurons (Figure 2D-F). This suggests a potential synaptic functional deficit in iNeurons lacking *SCN2A*, with a stronger effect in homozygous KO iNeurons. The G1744* iNeurons showed a complex phenotype, which may reflect the clinical presentation.

**Figure 2.**
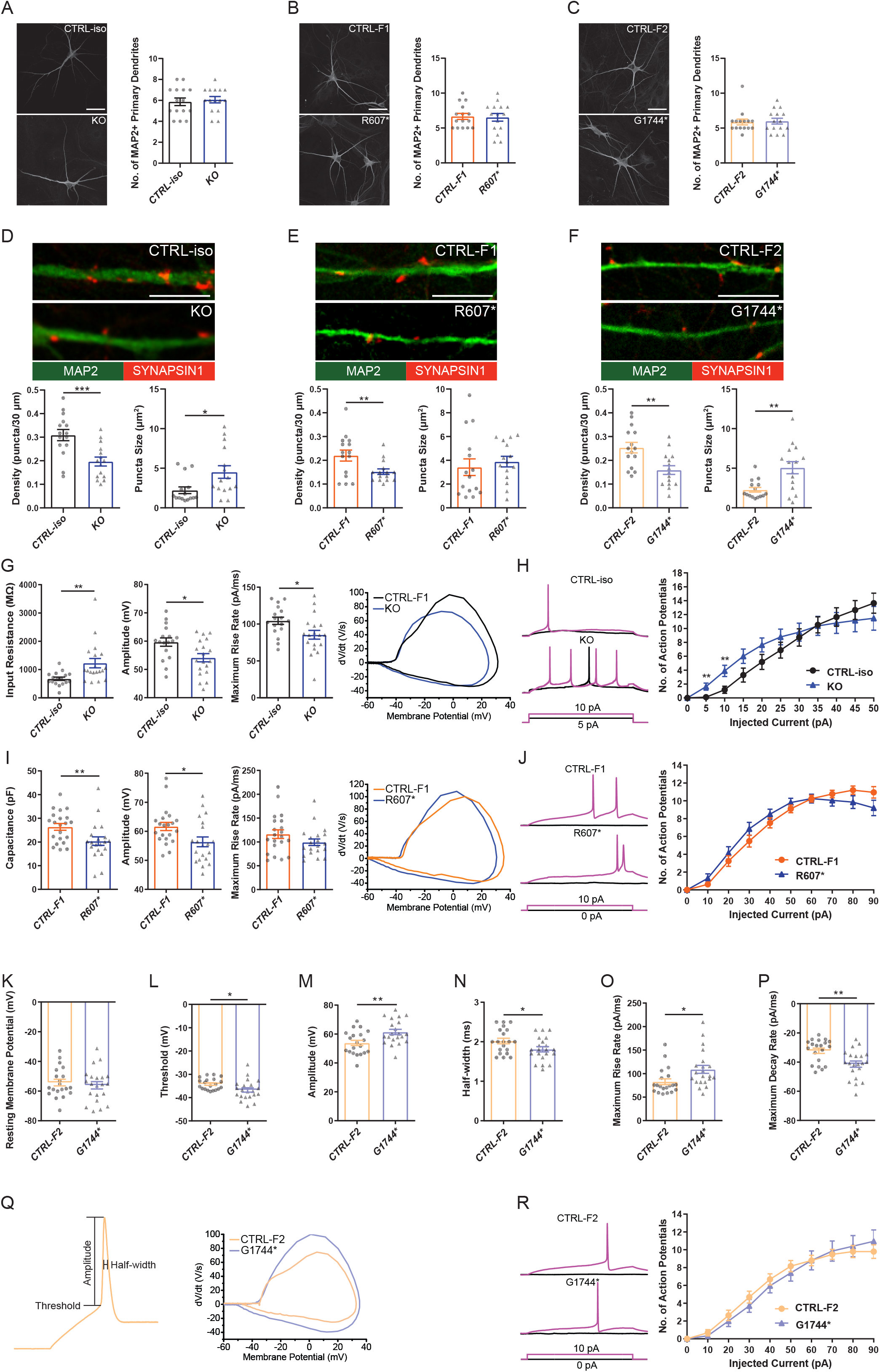
Effects of SCN2A *de novo* variants and CRISPR knockout on iNeuron morphology and synaptic function. See also figure S2. A-C) Representative images of immunocytochemistry for dendrite complexity and analysis of MAP2-positive primary dendrites. (A) CTRL-iso (n=15) and KO (n=15) iNeurons. (B) CTRL-F1 (n=15) and R607* (n=15) iNeurons. (C) CTRL-F2 (n=15) and G1744* (n=15) iNeurons. 3 viral transductions. Scale bar, 40 μm. D-F) Representative images and quantification of synaptic puncta density and size. (D) CTRL-iso (n=15) and KO (n=15) iNeurons. (E) CTRL-F1 (n=15) and R607* (n=15) iNeurons. (F) CTRL-F2 (n=15) and G1744* (n=15) iNeurons. 3 viral transductions. Scale bar, 10 μm. G) Membrane and action potential properties of CTRL-iso and SCN2A KO iNeurons (n=17 and 19, respectively), 3 viral transductions. Left: bar graph of recorded input resistance. Middle: bar graph of measured action potential amplitude. Right: bar graph of the maximum rise rate and the associated phase plane plot of action potential kinetics. Data represent means ± SEM. *p < 0.05, **p < 0.01, Student’s t-test. H) Repetitive firing properties of CTRL-iso and KO iNeurons. CTRL-iso (n=17) and KO (n=19) iNeurons, 3 viral transductions. Data represent means ± SEM. *p < 0.05, **p < 0.01, Student’s t-test. I) Membrane and action potential properties of CTRL-F1 and R607* iNeurons (n=21 and 20, respectively), 3 viral transductions. Left: bar graph of recorded capacitance. Middle: bar graph of measured action potential amplitude. Right: bar graph of the maximum rise rate and the associated phase plane plot of action potential kinetics. Data represent means ± SEM. *p < 0.05, **p < 0.01, Student’s t-test. J) Repetitive firing properties of CTRL-F1 and R607* iNeurons. CTRL-F1 (n=21) and R607* (n=20) iNeurons, 3 viral transductions. Data represent means ± SEM. *p < 0.05, Student’s t-test. K-Q) Membrane and action potential properties of CTRL-F2 and G1744* iNeurons (n=20 and 21, respectively), 3 viral transductions. (K) resting membrane potential, (L) threshold of the action potential, (M) amplitude of the action potential relative to threshold, (N) half-width of the action potential, (O) maximum rise rate from action potential threshold to amplitude, (P) maximum decay rate from action potential amplitude to threshold, (Q) representative action potential depicting measurements for analysis and the associated phase-plane plot of the respective action potential kinetics. R) Repetitive firing properties of CTRL-F2 and G1744* iNeurons. CTRL-F2 (n=20) and G1744* (n=21) iNeurons, 3 viral transductions. Data represent means ± SEM. *p < 0.05, Student’s t-test.

We next examined the biophysical properties of isogenic KO and patient-derived iNeurons. Whole-cell patch-clamp recordings were done at DIV 24 – 28, with all iNeurons being co-cultured with mouse glia to promote maturation and synapse formation. We measured intrinsic membrane properties, including active and passive electrophysiological properties. iNeurons were held at -70 mV membrane potential before evoking action potentials. We used hyperpolarizing currents to measure input membrane resistance.

KO iNeurons expressed higher input resistances compared to their isogenic controls (Figure 2G), suggesting that KO iNeurons require less current input to elicit the same response as CTRL-iso iNeurons. We also found KO iNeurons had decreased action potential amplitudes (Figure 2G), however, other intrinsic properties remained unchanged (Figure S2A-D). We analyzed action potential characteristics and found KO iNeurons had reduced maximum rise rates, (Figure 2G) which was similar to previous mouse studies (Spratt et al., 2021). There was no difference in the maximum decay rates or the half-width, suggesting that sodium channels were solely impaired (Figure S2E and F). To depict changes of action potential waveforms, we used phase-plane plots generated from the first derivative of the membrane potential (dV/dt) versus the membrane potential. This visualization depicts decreases in peak amplitude and the maximum value of dV/dt (Figure 2G). Next, we examined repetitive firing using a step protocol and found increases in the maximum number action potentials fired at 5 pA and 10 pA current injection steps for KO iNeurons (Figure 2H). This finding indicates a mild hyperexcitability phenotype. While this is paradoxical given that sodium channels are required for action potential generation, this phenomena is observed in conditional *Scn2a*^*-/-*^ mice (Spratt et al., 2021; Zhang et al., 2021).

We examined patient-derived iNeurons expressing the *de novo* R607* and G1744* truncating variants, and compared to their respective familial controls. Passive and active membrane properties were examined, and we determined the capacitance of R607* iNeurons was decreased relative to CTRL-F1 iNeurons (Figure 2I). This suggests that R607* iNeurons may be smaller, but other membrane properties remained unchanged (Figure S2G-I). However, we did not observe any changes to the resting membrane potential (Figure 2K) and other membrane properties in G1744* iNeurons (Figure S2M and N), suggesting this mutation does not affect cell size. When we measured action potential characteristics, we found an observable decrease in action potential amplitude in R607* iNeurons (Figure 2I), similar to isogenic KO iNeurons. However, maximum rise rate (Figure 2I) and other action potential characteristics remained unchanged in patient R607* iNeurons (Figure S2J-L), which distinguished this line from the KO. Measurement of the repetitive firing capabilities of R607* iNeurons revealed no statistical differences (Figure 2J), whereas there was a difference in isogenic KO iNeurons (Figure 2H), suggesting a milder phenotype. Analysis of G1744* iNeurons, however, showed stark differences compared to R607*. G1744* iNeurons had a shift in the action potential threshold (Figure 2L), which suggests the neurons could elicit an action potential at a more hyperpolarized voltage, a phenotype not observed in KO or R607* iNeurons. Furthermore, G1744* iNeurons produced more robust action potentials by measurement of their peak amplitude (Figure 2M). The action potential shape had a reduced half-width, suggesting that action potentials were narrower (Figure 2N). The altered action potential shape was supported by a significant increase in the rise and decay rate of the up and down stroke of the action potential relative to the threshold (Figure 2O and P). This is also visualized in the phase-plane plot (Figure 2Q). Together, these data reveal that G1744* iNeurons have faster action potential dynamics compared to KO and R607* iNeurons, demonstrating key differences between the variants. Lastly, repetitive firing capabilities of G1744* iNeurons revealed found no statistical differences (Figure 2R), similar to R607* iNeurons (Figure 2J). These data support the notion that SCN2A truncating variants can produce a spectrum of cellular phenotypes.

### Early truncation of SCN2A impairs excitatory synaptic transmission

We recorded spontaneous excitatory postsynaptic currents (sEPSCs) as a proxy for synaptic transmission to understand potential differences between the SCN2A variants. Isogenic KO iNeurons displayed a reduction in the frequency but not the amplitude of synaptic events (Figure 3A), similar to KO iNeurons derived from a different genetic background that we previously reported (Deneault et al., 2018). Recordings of R607* iNeurons revealed a similar decrease in sEPSC frequency with no change in amplitude (Figure 3B). However, there were no differences in sEPSC frequency or amplitude in G1744* neurons (Figure 3C). These experiments reveal that the KO and the R607* variants are largely loss-of-function and cause a reduction in synaptic connectivity and transmission, where the G1744* variant does not affect synaptic transmission but rather impacts action potential properties. These data indicate that the C-terminal G1744* variant is not a typical loss of function protein-truncating variant, suggesting that the nonsense-mediated decay site may be important for determining the type of channel impairment in addition to the variant location after the breakpoint.

**Figure 3.**
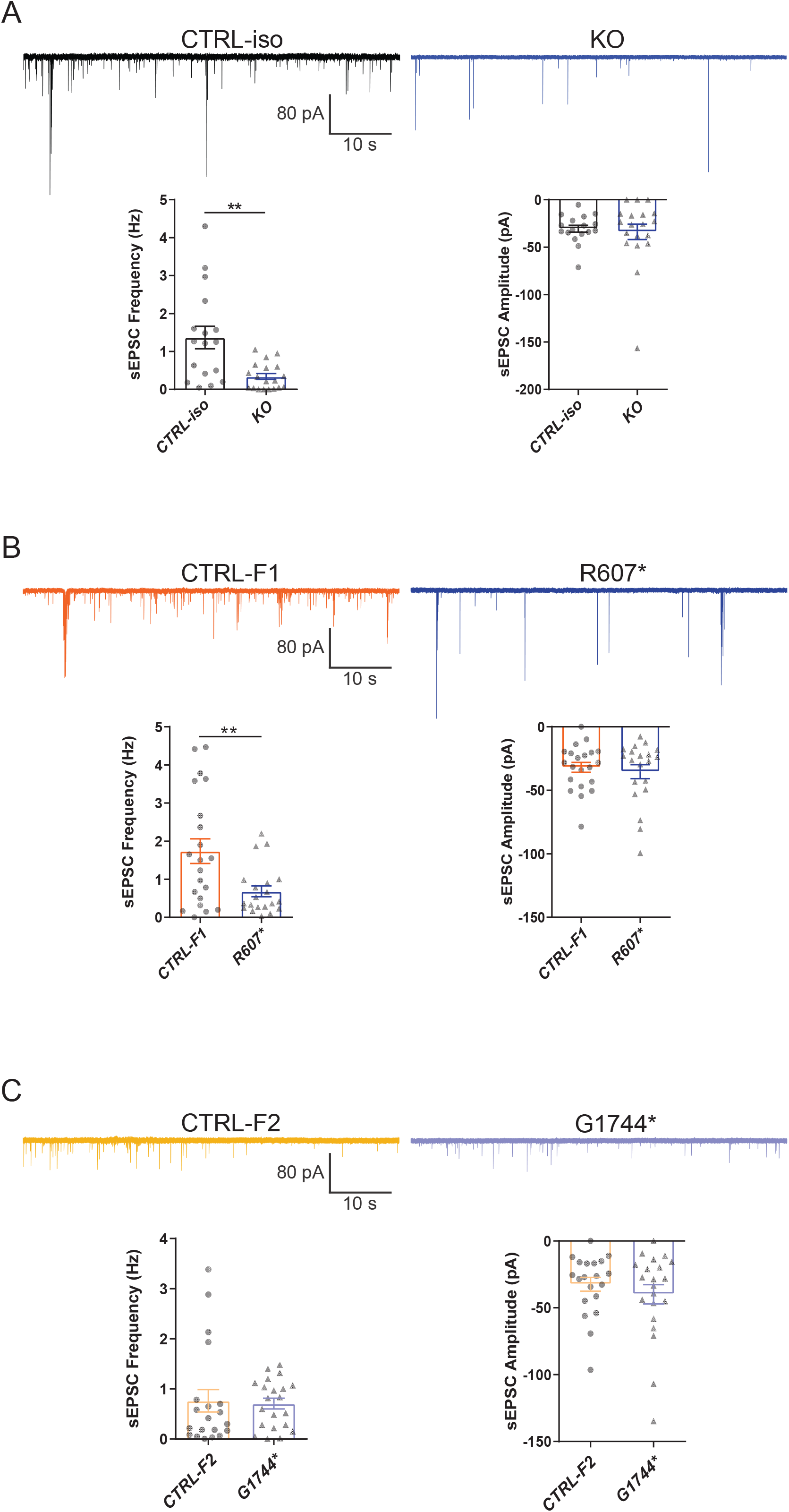
Complete and partial loss of SCN2A has differential effects on synaptic transmission. A-C) Synaptic transmission representative traces and analysis of SCN2A deficient iNeurons. (A) Left: sEPSC frequency of synaptic transmission. Right: sEPSC amplitude of synaptic transmission. CTRL-iso (n=17) and KO (n=19) iNeurons. (B) Left: sEPSC frequency of synaptic transmission. Right: sEPSC amplitude of synaptic transmission. CTRL-F1 (n=21) and R607* (n=20) iNeurons. (C) Left: sEPSC frequency of synaptic transmission. Right: sEPSC amplitude of synaptic transmission. CTRL-F2 (n=20) and G1744* (n=21) iNeurons. 3 viral transductions. Data represent means ± SEM. *p < 0.05, **p < 0.01, Student’s t-test.

### Severe loss of SCN2A impairs spontaneous neuronal network activity while the G1744* variant shows early transient impairments in iNeurons

To better understanding how disruption of *SCN2A* can affect the longitudinal development of neuronal circuits in a network, we used microelectrode arrays (MEAs). Cultures followed the same plating protocols (Figure 1A), but were optimized for long-term survival (> 4 weeks *in vitro*). Isogenic KO, R607* raster plots were taken from DIV 49 depicting network activity within a well, highlighting the activity differences versus the respected control, and similarities and differences between the genotypes; G1744* iNeuron raster plots were taken from DIV 28 since phenotypes were strongest during early development and were transient (Figure 4A, H and O).

**Figure 4.**
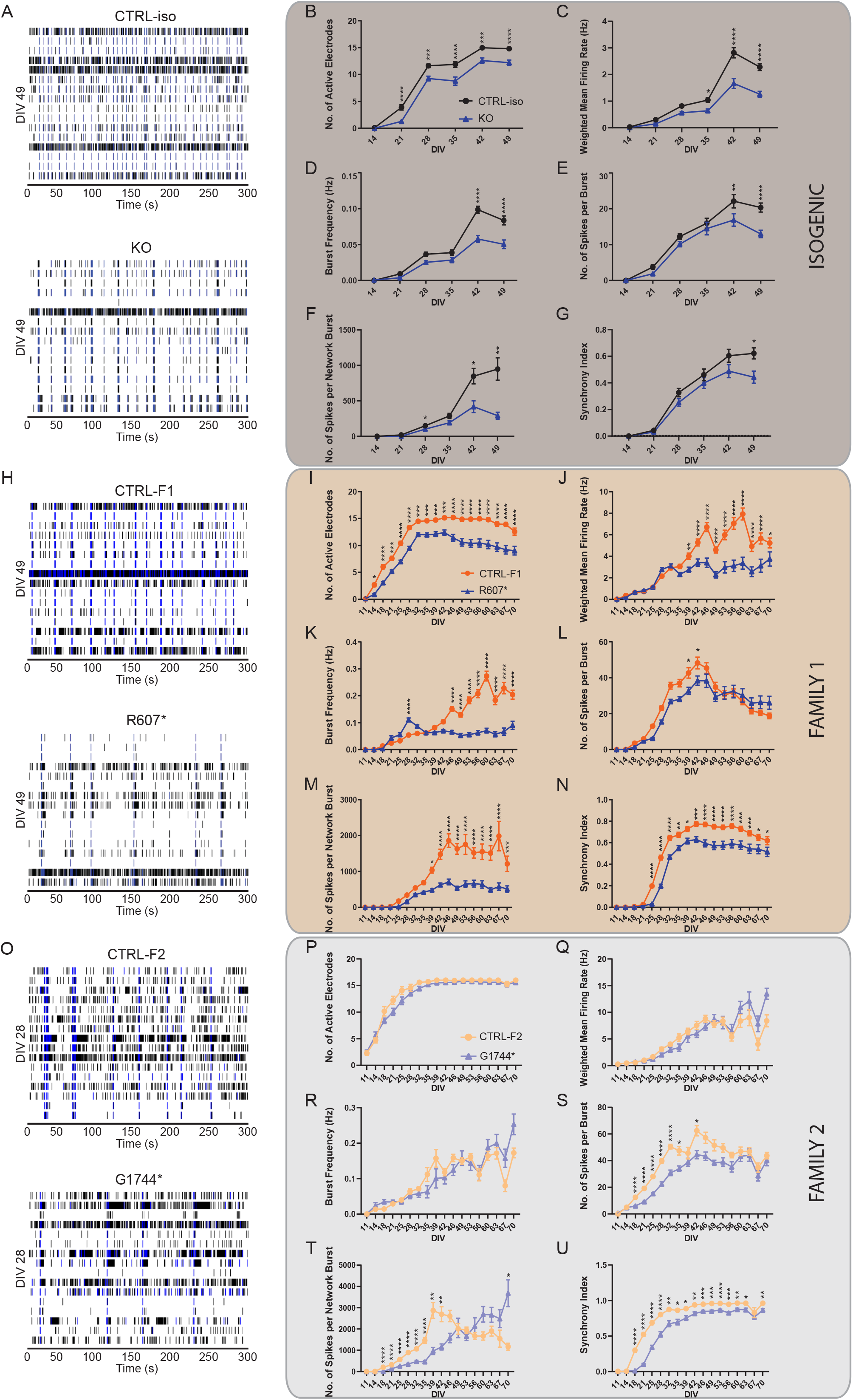
Effects of SCN2A *de novo* variants and knockout on the development of spontaneous network activity in iPSC-derived iNeurons. See also figure S3. A) Example raster plots of recordings of neuronal network activity at DIV 49 for CTRL-iso and SCN2A KO iNeurons. B-G) Quantification of MEA parameters for CTRL-iso and KO iNeurons. (B) number of active electrodes, (C) weighted mean firing rate, (D) burst frequency, (E) number of spikes per burst, (F) number of spikes per network burst, (G) synchrony index. CTRL-iso (n=47 wells) and KO (n=43 wells) iNeurons, 3 viral transductions. Data represent means ± SEM. *p < 0.05, **p < 0.01, ***p < 0.001, ****p < 0.0001, two-way ANOVA with post hoc Sidak correction. H) Example raster plots of recordings of neuronal network activity at DIV 49 for CTRL-F1 and R607* iNeurons. I-N) Quantification of MEA parameters for CTRL-F1 and R607* iNeurons. (I) number of active electrodes, (J) weighted mean firing rate, (K) burst frequency, (L) number of spikes per burst, (M) number of spikes per network burst, (N) synchrony index. CTRL-F1 (n=48 wells) and R607* (n=46 wells) iNeurons, 3 viral transductions. Data represent means ± SEM. *p < 0.05, **p < 0.01, ***p < 0.001, ****p < 0.0001, two-way ANOVA with post hoc Sidak correction. (O) Example raster plots of recordings of neural network activity at DIV 28 for G1744* and CTRL-F2 iNeurons. P-U) Quantification of MEA parameters for CTRL-F2 and G1744* iNeurons. (P) number of active electrodes, (Q) weighted mean firing rate, (R) burst frequency, (S) number of spikes per burst, (T) number of spikes per network burst, (U) synchrony index. CTRL-F2 (n=20 wells) and G1744* (n=19 wells), 2 viral transductions. Data represent means ± SEM. *p < 0.05, **p < 0.01, ***p < 0.001, ****p < 0.0001, two-way ANOVA with post hoc Sidak correction.

Overall, both SCN2A KO and R607* iNeurons displayed similar trends in various parameters. Both had a decrease in active electrodes across development (Figure 4B and I), which was not due to reduced neuronal survival (data not shown). Examining the weighted mean firing rate (wMFR), which accounts for the decrease in active electrodes, revealed a reduction in KO and R607* iNeurons that began early and persisted (Figure 4C and J). This indicates that the loss of *SCN2A* hinders the abundance of neuronal spikes within the network. We extended the experiment time of patient neurons up to 10-weeks to determine if longer-term culture conditions alter their developmental trajectory. We found that up to 10-weeks, there was still an overall decrease in R607* iNeuron firing. G1744* iNeurons on the other hand did not show any remarkable deficits in active electrodes or wMFR across the 10-week time point, demonstrating no major firing differences by this variant (Figure 4P and Q).

We further investigated whether network bursting was disrupted since this parameter is indicative of population neuronal activity, and it is disrupted in other models of SCN2A deficiency (Deneault et al., 2018; Lu et al., 2019). Additionally, recent investigations into risk genes contributing to ASD and intellectual disability clinical features have showed changes in neuronal network activity, specifically enhancements of bursts (Frega et al., 2020). We examined burst frequency and the number of spikes within each burst since sodium channels are important in action potential and spike generation. Bursts were defined by having 5 spikes with a maximum of 100 millisecond inter-spike intervals. KO iNeurons displayed a decrease in bursting and the number of spikes per burst (Figure 4D, E). When examining the bursting of patient R607* iNeurons, unlike the KO iNeurons, R607* iNeurons exhibited a small increase in bursting early in development (Figure 4K), which dissipated and became reduced compared to CTRL-F1 iNeurons. Furthermore R607* iNeurons displayed a decrease in the number of spikes per burst for a transient window between DIV 39 – 46 (Figure 4E vs. L). Secondary burst parameters were also examined to further characterize bursting patterns (Figure S3A-B and E-F). Interestingly, G1744* iNeurons did not show any difference in burst frequency but had a stark reduction in the number of spikes per burst during early development (Figure 4R and S, Figure S3I and J).

Network bursting and synchronization occurs in later stages of neuronal development, where this is crucial for the organization and regulation of excitation (Kim et al., 2019; Kirwan et al., 2015; Masquelier and Deco, 2013; Stegenga et al., 2008). Network bursts were defined as having 4 or more electrodes to burst within 100 ms of one another. We found that the number of spikes per network burst was reduced in KO and R607* iNeurons at approximately 4 weeks (Figure 3F and M). Network burst frequency and network burst duration were also recorded and corroborated impaired network capacity (Figure S3C-D and G-H,). Interestingly, we found that early on within network bursts, G1744* iNeurons had significantly reduced spikes, similar to KO and R607* iNeurons; however, after DIV 49, there was a reversal and G1744* iNeurons had significantly more spikes (Figure 4T). This was also reflected in the large increase in the burst and network burst duration (Figure S3K and L), which is opposite to the KO and R607* iNeurons. This is noteworthy because this proband has early-onset seizure activity; therefore, the increased spikes per network burst and their duration, which may occur due to the abnormal action potential characteristics, may contribute to the epilepsy. An alternative explanation is that the G1744* iNeurons have an elongated early maturation phase where normal developing burst sequences are altered initially, but compensate for the decreased number of spikes in bursts allowing for a rebound in mid to late development.

Lastly, we examined synchronization, which is a parameter defined as the probability for neighboring electrodes to detect activity in quick succession based on prior electrode activity. This is measured as an index ranging from 0 – 1, where 1 represents neighboring electrodes detecting activity based on a prior active electrode, 100% of the time. Interestingly, we found that all iNeurons displayed deviation from their respected control lines, suggesting that synchronization of neurons in the network was impaired by any type of SCN2A disruption (Figure 4G, N and U). However, this measurement does not take into account the probability of random spikes contributing to the synchrony of the neuronal network.

### Proteomic analysis of SCN2A R607* iNeurons reveals neuronal morphological and bioenergetic signaling deficits

To better understand the molecular impact of the patient *SCN2A* variants in iNeurons, we performed quantitative TMT-tagged shotgun proteomics at DIV 14 iNeurons focusing on the R607* variant since it had the more severe phenotypes compared to G1744*. We used a single batch experiment method with 3 biological replicates to reduce variability and plated iNeurons without glia to avoid cross identification with mouse glial proteins. Similar plating methods have been used previously for RNA sequencing from iNeurons (Deneault et al., 2018). We TMT-labeled equal quantities of sample (see methods), and identified 7516 proteins with at least two unique peptides for Family 1. At DIV 14, each condition clustered together as shown by principal component analysis plot (Figure 5A). We found 2337 significantly differentially expressed proteins (p-adjusted < 0.05, FC > 1.2 or FC < 0.8) (Figure 5B). We used g:Profiler for functional enrichment analysis (Figure 5C) using both increased and decreased differentially expressed proteins (Reimand et al., 2016, 2019). Among the enrichment clusters were Mitochondrial Function, Neuron Projection Development, Translation, and Cytoskeletal Organization (Figure 5C and S4). Further examination of the enriched clusters for the GO:Biological process terms revealed changes in Mitochondrial function: organization and translation, and Neuron projection development: neuron development and differentiation (Figure 5D). The differentially expressed proteins, both upregulated and downregulated belonging to each of the 2 clusters of interest are shown (Figure 5E). Of note, we found a reduction in expression of key mitochondrial proteins such as TOMM20 (p-adj = 0.00489, Log2FC = -1.05) and the mitochondrial fusion regulators Mitofusin 1 (MFN1) (p-adj = 0.002, Log2FC = -0.34) and Mitofusin 2 (MFN2) (p-adj = 0.004, Log2FC = -0.487). These data suggest R607* neurons may have altered mitochondria numbers or functionality (Chen et al., 2003; Filadi et al., 2015, 2018; Pecorelli et al., 2020).

**Figure 5.**
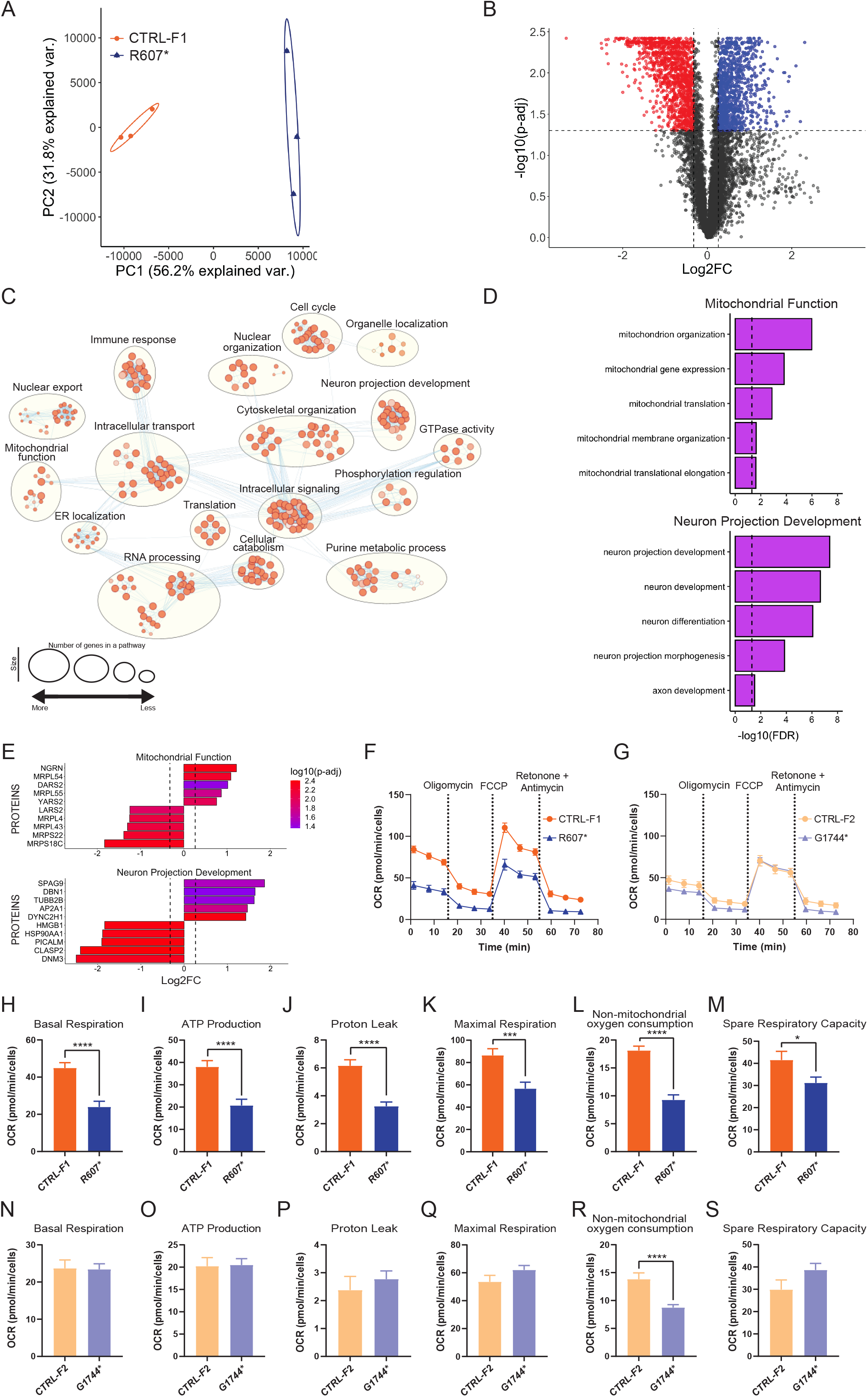
Proteomic analysis of CTRL-F1 and SCN2A R607* iNeurons. See also figure S4. A) PCA plot showing clustering of the 3 biological replicates for each genotype used in proteomics. B) Volcano plot showing differentially expressed proteins. Downregulated proteins are shown in red and upregulated proteins are shown in blue. Gray points are non-differentially expressed proteins. Horizontal dashed line represents p-adjusted threshold of 0.05 and vertical dashed lines represent threshold of ± 20% foldchange on a log_2_ scale. C) Network visualization of g:Profiler enrichment of GO:Biological Processes (BP) of both increased and decreased DEPs. Orange dots represent over-represented GO:BP terms. A large circle indicates the number of genes within a term and darker shades represent a higher degree of significance found during enrichment. Clusters indicate biological pathways with common proteins and function. D) GO analysis of clusters from C) showing selected GO:BP terms. E) Top differentially expressed proteins from clusters identified in C). F-G) Live-cell metabolic assay for validation studies of R607* iNeuron proteomics. (F) Line graph depicting changes in R607* and CTRL-F1 iNeurons in Seahorse assay. (G) Line graph depicting G1744* and CTRL-F2 iNeurons in Seahorse assay. H-M) Analysis of mitochondrial respiration phases for R607* and CTRL-F1 iNeurons. (H) basal respiration, (I) ATP production, (J) proton leak from mitochondria, (K) maximal respiration from mitochondria, (L) non-mitochondrial oxygen consumption, (M) spare respiratory capacity. CTRL-F1 (n=45 wells) and R607* (n=45 wells), 3 viral transductions recorded at DIV 14. Data represent means ± SEM. *p < 0.05, **p < 0.01, ***p < 0.001, ****p < 0.0001, Student’s t-test. N-S) Analysis of mitochondrial respiration phases for G1744* and CTRL-F2 iNeurons. (N) basal respiration, (O) ATP production, (P) proton leak from mitochondria, (Q) maximal respiration from mitochondria, (R) non-mitochondrial oxygen consumption, (S) spare respiratory capacity. CTRL-F2 (n=45 wells) and G1744* (n=45 wells), 3 viral transductions recorded at DIV 14. Data represent means ± SEM. *p < 0.05, **p < 0.01, ***p < 0.001, ****p < 0.0001, Student’s t-test.

We hypothesized the potential changes in mitochondrial function and bioenergetic signaling could be contributing to R607* synaptic phenotypes. Further, metabolism, NDDs, and neuronal dysfunction have been reported to be highly interconnected (Ebrahimi-Fakhari et al., 2016; Lewis et al., 2018; Li et al., 2019; Rangaraju et al., 2019; Rose et al., 2018), including a close relationship between mitochondrial function and neuronal firing (Ruggiero et al., 2021). To functionally validate this pathway, we used iNeurons (co-cultured with mouse glial cells) as they have been previously used to study mitochondrial dysfunction (Li et al., 2019). We used a Seahorse XF96 analyzer with the mitochondrial stress test (MST) protocol. This assay measures mitochondrial respiration as oxygen consumption rate (OCR), and the addition of 4 compounds (Oligomycin, carbonyl cyanide-4-(trifluromethoxy)phenylhydrazone (FCCP), and Rotenone/Antimycin A), to disrupt the electron transport chain (ETC) and analyze the contributions of various components. Three independent transductions of CTRL-F1 and R607* iNeurons were co-cultured with mouse glia in a 96-well plate format and analyzed at DIV 14. We found significantly altered mitochondrial respiration in patient R607* iNeurons (Figure 5F). We found that R607* iNeurons had reduced baseline respiration compared to CTRL-F1 (Figure 5H), as well as reduced adenosine triphosphate (ATP) production, proton leak, maximal respiration, non-mitochondrial oxygen consumption and spare respiratory capacity (Figure 5I-M). Decreases in ATP production was supported by components of mitochondrial ATP synthase subunit 5 being downregulated (ATP5F1A p-adj = 0.033, Log2FC = -0.461 and ATP5F1E p-adj = 0.011, Log2FC = -0.340), while other mitochondrial respiration deficits were supported by downregulated proteins necessary for mitochondrial activity. We also examined mitochondrial function in G1744* iNeurons given the differences in synaptic function compared to R607* iNeurons (Figure 5G). We found that a small change in mitochondrial function in G1744* iNeurons, supporting the idea that the G1744* variant does not impact synaptic transmission and has a different phenotype that R607* (Figure 5N-S). Decreases in the non-mitochondrial oxygen consumption may suggest that G1744* are less energetic or have a slower metabolism. Taken together, R607* expressing iNeurons potentially have a disruption of mitochondrial respiration and homeostasis compared to CTRL-F1, suggesting that altered bioenergetics and mitochondrial/metabolic function maybe the downstream results of the loss of SCN2A function.

## DISCUSSION

In this study, we present data from three genetic human cellular models of *SCN2A*, to understand the channelopathy contributions to neurodevelopmental disorders. Models suggest that *SCN2A* variants that cause an enhancement of neuronal excitability result in epilepsy and are gain-of-function, whereas, ASD-linked variants are loss-of-function and reduce channel activity (Begemann et al., 2019; Ben-Shalom et al., 2017; Deneault et al., 2018; Lu et al., 2019; Ogiwara et al., 2018; Rubinstein et al., 2018; Sanders et al., 2018; Spratt et al., 2021, 2019; Wang et al., 2021; Zhang et al., 2021). This classification does not fully characterize the channelopathy, especially where an estimated 20 – 30% of ASD/ID patients develop late-onset seizures in addition to SCN2A deficiency (Sanders et al., 2018; Wolff et al., 2017; Zhang et al., 2021). We found that patient iNeurons harboring different ASD-linked SCN2A variants do not have the same phenotype, but exhibit differences in action potential characteristics, synaptic dysfunction and neuronal signaling networks. This indicates that the non-sense mediated decay breakpoint could play a role in determining the functional impact of an SCN2A truncating variant.

Human iPSC-derived KO iNeurons displayed a paradoxical hyperexcitability phenotype, similar to mouse Scn2a^-/-^ neurons (Spratt et al., 2021; Zhang et al., 2021), but not in patient iNeurons as they have a heterozygous loss of SCN2A function. The reduction in synaptic transmission in KO iNeurons could be due to the localization of SCN2A during early development predominately being expressed at the axon initial segment and secondarily in the somatodendritic compartments (Bender and Trussell, 2012; Gazina et al., 2015; Hu and Bean, 2018; Hu et al., 2009; Spratt et al., 2019). An alternative explanation is that SCN2A regulates neuronal activity-dependent transcriptional activity that drives synaptic gene expression, and protein translation which was disrupted in our proteomics analysis (Figure S4) (Deneault et al., 2018; Ip et al., 2018; Madabhushi and Kim, 2018; Nelson and Valakh, 2015; Zhang et al., 2018).

When comparing the R607* to the G1744* variant, R607* had a similar phenotype to KO neurons than G1744* (except the hyperexcitability). R607* iNeurons had a decrease in synaptic transmission, but it is unknown if this is a direct or indirect impact. Extrapolating and comparing to mouse Scn2a haploinsufficiency, the reduction in excitatory synaptic transmission in human R607* iNeurons could be driven by changes in the AMPA:NMDA ratio, inferring an abundance of silent synapses due to an abundance of AMPA-lacking spines (Hanse et al., 2013; Kerchner and Nicoll, 2008; Spratt et al., 2019). We noted both KO and R607* iNeurons generated less spontaneous electrical activity, which could suggest SCN2A deficiency, without seizures, dampens neural networks and produces small quiescent pockets within the network (Deneault et al., 2018). Future studies can examine how other SCN2A missense variants impact human neuron synaptic function, as these variants do not cause protein truncation, but only amino acid changes, leading to effects on channel function (Ben-Shalom et al., 2017; Echevarria-Cooper et al., 2021).

The G1744* variant is loss-of-function but the individual presents with early-onset seizures, and the variant is past the predicted non-sense mediated decay breakpoint (Nagy and Maquat, 1998; Sanders et al., 2018). The major finding from this line was an enhancement of the action potential waveform, similar to a human neuronal model of the SCN2A-L1342P variant (Que et al., 2021). However, the L1342P variant is gain-of-function from computational modelling and its association with epilepsy, but it has mixed gain- and loss-of-function phenotypes (Begemann et al., 2019; Que et al., 2021). Even with action potential waveform enhancements, we did not observe increases in neuronal excitability, dissimilar to the L1342P variant. Unlike previous SCN2A mouse and human studies, we did not observe a difference in synaptic transmission of G1744* iNeurons (Deneault et al., 2018; Spratt et al., 2019; Wang et al., 2021), suggesting there is compensation for the reduction in synapse formation. The differences between R607* and G1744* underscore the importance of studying multiple variants in sodium channels to ascertain the impact on function, as this was shown for *SCN8A* (Lopez-Santiago et al., 2017). In G1744* iNeurons, we speculate the reduction of spikes during early development was compensated by the early onset seizures. This could explain why at later timepoints the only difference was the synchronization of the network with no bursting (Que et al., 2021; Tidball et al., 2020). Additionally, G1744* iNeurons may have compensatory activity early in development and recover, which may be due to a truncated protein being produced that avoids non-sense mediated decay. Given that the G1744 position is near the calmodulin-binding domain of SCN2A, variants near the C-terminus may impact local calcium signaling, which has yet to be examined.

The proteomic profiling revealed that the R607* patient iNeurons had significantly impaired neuronal development, translation and synaptic signaling networks. The validation that mitochondrial and metabolic function is defective in R607* neurons suggests that SCN2A, neuronal activity and mitochondrial activity are coupled. Mitochondrial dysfunction has previously been associated with ASD and NDDs (Frye, 2020; Rossignol and Frye, 2012) and up to 80% of ASD cases have biomarkers of mitochondrial and ETC activity dysfunction (Rossignol and Frye, 2012). Recent reports have also linked mitochondrial dynamics (fission and fusion) to altered by neuronal activity (Divakaruni et al., 2018; Lee et al., 2018; Rangaraju et al., 2019; Sung et al., 2008). This could link how the reduction in synaptic activity of the R607* variant leads to impaired mitochondrial function, which impacts ATP and energy stores, the reduced mitochondrial respiration could also be a key driver of synaptic deficits acting as a positive feedback loop in R607* iNeurons. Several of the measured parameters showed that mitochondria are not functioning appropriately. Mitochondrial function has also been implicated in specific genetic forms of ASD or NDDs (Cicaloni et al., 2020; Crivellari et al., 2021; Ebrahimi-Fakhari et al., 2016; Gebara et al., 2021; Klein Gunnewiek et al., 2020; Kwan et al., 2020; Mithal and Chandel, 2020), suggesting convergence on this pathway is a risk factor.

The NGN2 model system has both advantages and disadvantages for modeling *SCN2A* variants. The system is robust in producing functional neurons from iPSCs, and iNeurons function well for in multielectrode array and patch-clamp experiments. However, the limitations are that iNeurons are more heterogeneous than previously thought, and do not resemble a specific neuronal sub-type (Lin et al., 2021). Further, their transcriptional signatures can be shared with sensory neurons (Lin et al., 2021), and linked to NGN2 expression levels. While we addressed this to some degree by using the same viral titer (and infection protocol), this may not be sufficient. Further, a small portion of iNeurons can have two or more axons (Rhee et al., 2019). Therefore, future experiments on *SCN2A* variants should be tested in both the iNeuron and NPC-neuron systems.

Taken together, our results indicate that *SCN2A* variants can have different functional impacts on human neurons, and variants associated with ASD and seizures may not possess a typical loss-of-function effect. Variants with the greatest impact on SCN2A expression seemed to have the biggest impact on core neuronal activity, but this led to more complex downstream phenotypes. Further analysis of different *SCN2A* variants in neurons, across the gene, will be required to understand the functional impact, and to better predict the potential effects on circuit dysfunction in the brain.

## ACKNOWLEDGEMENTS

Funding for the study was provided by grants from the Canadian Institutes of Health Research (CIHR), Ontario Brain Institute-POND study and the Natural Sciences and Engineering Research Council (NSERC) to K.K.S. Y.L received funding from NSERC and ERA-NET NEURON, and S.W.S received funding from OBI-POND, Autism Speaks and CIHR. J.U was awarded a fellowship from CIHR (CGS-M) and the University of Toronto Vision Science Research Program, and C.O.B was awarded a fellowship from the McMaster University Michael G. DeGroote Institute for Pain Research and Care. The authors wish to acknowledge the resources of MSSNG (www.mss.ng), Autism Speaks and The Centre for Applied Genomics at The Hospital for Sick Children, Toronto, Canada. We also thank the participating families for their time and contributions to this database.

## AUTHOR CONTRIBUTIONS

C.O.B and K.K.S contributed to experimental conception and planning. C.O.B contributed to the majority of experimentation, writing and editing of the manuscript. J.U contributed to experimentation and writing. E.R, A.A, S.X, B.M.D, S.K, S.H.W and Y.L contributed to experimentation. A.A.C contributed to the generation of resources. J.H coordinated sample collections. N.M, Y.L, S.W.S and K.K.S contributed to conceptual development and review of the manuscript. K.K.S supervised C.O.B, N.M, J.U and A.A.C.

## EXPERIMENTAL PROCEDURES

### Approval and generation of iPSCs

All pluripotent stem cell work was approved by the Canadian Institutes of Health Research Stem Cell Oversight Committee. Blood was taken from individuals with the approval from SickKids Research Ethics Board after informed consent was obtained, REB approval file 1000050639. This study was also approved by the Hamilton Integrated Research Ethics Board, REB approval file #2707. CD34+ blood cells were verified using flow cytometry and collected for iPSC reprogramming. All iPSCs were generated by Sendai virus reprogramming and clonal expansion using the CytoTune – iPSC 2.0 kit (ThermoFisher) to deliver the reprogramming factors. Once colonies were large enough (approximately 15-17 days post Sendai transduction), each colony was transferred to 1 well of a 12-well plate coated with irradiated MEFs and plated in iPSC media (DMEM/F12 supplemented with 10% KO serum, 1x non-essential amino acids, 1x GlutaMAX, 1mM β-mercaptoethanol, and 16ng/mL basic fibroblast growth factor (bFGF)). Once iPSCs were expanded and established they were transitioned to matrigel coated plates and grown in mTeSR1 (STEMCELL Technologies) and subsequent passaging continued to use ReLeSR (STEMCELL Technologies). iPSC lines without karyotypic abnormalities were used and this was verified by G-band karyotyping performed by the Centre for Applied Genomics (Hospital for Sick Children). To verify the expression of pluripotent markers OCT4 and NANOG, immunocytochemistry was performed.

### Generation of *SCN2A* KO cells

A 3x stop premature termination codon (StopTag) was designed similarly to our previously described method for introducing a StopTag into the DNA to knock out the expression of genes of interest (Deneault et al., 2018). This 3x StopTag was delivered by a synthesized single-stranded oligodeoxynucleotide (ssODN) containing two 50-nucleotide-long homology arms with the StopTag, V5 epitope and EcoRI restriction site sequences coding 59 nucleotides. The knock in site was selected to situate the StopTag 80 bp upstream of the exon 5-6 junction, to increase the likelihood of transcript knockout by the nonsense-mediated decay pathway (Neu-Yilik et al., 2011). A normal iPSC line named “50B” previously described was used to insert the StopTag and generate an isogenic *SCN2A* KO line (Deneault et al., 2018). The StopTag ssODN template, a pSpCas9(BB)-2A-GFP plasmid (Addgene, catalog no. 48138) and paired gRNAs targeting exon 5 of *SCN2A* were nucleofected into the 50B iPSCs using the Amaxa 4D-nucleofector with code CA137. GFP expressed cells were isolated 48h after nucleofection and clonally grown. Digital droplet PCR and dilution culture steps previously described were used to enrich for *SCN2A* KO populations (Deneault et al., 2018; Miyaoka et al., 2014). The purified *SCN2A* KO wells were expanded and assayed for pluripotency, karyotypic abnormalities and sequencing validation of StopTag insertion. (Figure S1).

### Induction of iPSCs into glutamatergic neurons

We sought to explore functional differences of the 3 genetics models of *SCN2A* deficiency. For this, we needed to differentiate the newly generated iPSCs into excitatory neurons. Since previous findings showed inhibitory neurons were unaffected by *SCN2A* deficiency, we required an established system to explore excitatory neuron driven differences (Ogiwara et al., 2018; Spratt et al., 2019; Wang et al., 2021). In order to rapidly upscale experiments and focus on excitatory neurons, we used the previously published constitutive expression protocol of NGN2 to generate homogeneous populations of glutamatergic neurons. (Zhang et al., 2013). These iNeurons displayed stable membrane, firing and synaptic properties when co-cultured with mouse glial cells by DIV 21 (Zhang et al., 2013, 2018). Importantly as we previously described, this protocol provided consistent differentiation levels between cell lines derived from different participants (Deneault et al., 2018, 2019). We modified this protocol by inducing NGN2 for 3 days starting at DIV 1 and puromycin selecting for 2 days starting at DIV 2 and adding mouse glial cells at DIV 5. Half-iNeuron media (iNI) (Neurobasal media, 1x SM1, 1x GlutaMAX 1x pen/strep, 1µg/mL laminin, 10 ng/uL BDNF and 10 ng/uL GDNF) changes were performed every other day. Patch-clamp recordings were generated between DIV 24 and 27 post-NGN2-induction. Comparable bioelectric properties were found to previously reported (Deneault et al., 2018, 2019; Yi et al., 2016; Zhang et al., 2013, 2018).

### Immunocytochemistry

iPSCs were washed gently 2 times with PBS and fixed in 4% paraformaldehyde in PBS for 8 min at room temperature. The cells were then washed 2 times with PBS and left overnight in 4°C. The next day, the cells were permeabilized in -30°C with ice cold methanol for 10 min. The cells were then washed 2 times with PBS for 8 min and incubated primary antibodies overnight at 4°C. The following day, the cells were washed 3 times with PBS for 8 min. Secondary antibodies were incubated for 1 hr at room temperature covered with aluminum foil, followed by 3 washes in PBS for 8 min. After the washes, 300 mM DAPI in PBS was incubated for 8 min, followed by 2 washes with PBS. Coverslips were then quicky dried with a Kimwipe, and mounted on VistaVision glass microscope slides (VWR) with 10 µL of Prolong Gold Anti-Fade mounting medium (Life Technologies). Mounted coverslips were allowed to cure overnight in a dark slide box at room temperature. Images were acquired using a Zeiss LSM700 confocal microscope.

On DIV 25, iNeurons were fixed at room temperature in 4% paraformaldehyde in PBS for 15 min. The cells were then washed 3 times for 10 min with PBS, then blocked and permeabilized (B/P) with a B/P solution containing (0.3% Triton-X, 10% Donkey Serum, and PBS) for 1 hr. The cells were then incubated overnight at 4°C with primary antibodies in B/P solution. The next day, cells were washed 3 times for 10 min in PBS and incubated with secondary antibodies in B/P solution for 1.5 hours at room temperature and covered with aluminum foil. The cells were then washed 3 times for 10 min and incubated with 300mM DAPI for 8 min. The cells were then washed 1 time with PBS for 10 min. Coverslips were then quicky dried with a Kimwipe, and mounted on VistaVision glass microscope slides (VWR) with 10 µL of Prolong Gold Anti-Fade mounting medium (Life Technologies). Mounted coverslips were allowed to cure overnight in a dark slide box at room temperature. Images were acquired using a Zeiss LSM700 confocal microscope.

Synaptic morphology was processed and analyzed with ImageJ software. The Synapsin1 antibody was co-immunostained with MAP2 to determine dendrites with presynaptic puncta. Three biological replicates were used for each line with the data generated from five iNeurons per replicate per condition. A total of 15 iNeurons per condition per line were used with two dendrites of equal dimensions used per iNeuron. Data represent the number of synaptic puncta averaged by two dendrites per iNeuron within 30 µm segments. The same images were used to calculate dendrite complexity. This was determined by counting the number of MAP2-positive primary dendrites branching from the soma.

### Multi-electrode array

All recordings were performed using 48-well clear bottom MEA plates (Axion Biosystems), consisting of 16 electrodes per well. Plates were coated with filter-sterilized 0.1% polyethyleneimine solution in borate buffer pH 8.4 for 2 hr at 37°C, washed with water four times and dried overnight. 40 000 DIV 4 doxycycline iNeurons were seeded in a 20 uL drop of iNI media at the centre of each well for 1.5 hr, then an additional 150 uL of iNI media was added. The day after, 20 000 mouse astrocytes per well were seeded on top of iNeurons in 150 uL per well of iNI media. Mouse astrocytes were prepared from postnatal day 1 CD-1 mice as described (Kim and Magrané, 2011). Half media changes were performed every other day with iNI media until the endpoint of experiments. The electrical activity of neurons was measured a minimum of once-a-week post-seeding onto MEA plates using the Axion Maestro MEA reader (Axion Biosystems). On the day of recording, MEA plates were equilibrated for 5 min on the pre-warmed reader at 37°C. Real-time spontaneous neural activity was recorded for 10 min to use for offline processing. Recordings were sampled at 10 kHz, and filtered with a bandpass filter from 200 Hz to 3 kHz. A threshold of greater than 6 standard deviations was used to detect spikes and separate noise. Electrodes were considered active if a minimum of 5 spikes were detected per minute. Wells that were unable to generate 10 active electrodes of the 16 by DIV 42 were not used for analysis. Bursts were defined as a minimum of 5 spikes with a maximum of 100 millisecond inter-spike interval (ISI). Network bursts were defined as a minimum of 10 spikes with a maximum of 100 milliseconds ISI and at least 35% of electrodes in synchrony. Offline processing was performed using Axion Biosystems Neural Metric Tool.

### *In vitro* electrophysiology

iNeurons were replated on DIV 4 onto polyornithine/laminin coated coverslips in a 24-well plate at a density of 100 000/well with 0.5 mL of iNI media. On DIV 5, primary mouse astrocytes were added at a density of 50 000/well to support iNeurons’ viability and maturation. Half media changes were performed every other day and wells were maintained until DIV 24 – 26 for recordings. At DIV 9, iNI was supplemented with 2.5% FBS which was adapted from (Zhang et al., 2013). Whole-cell patch-clamp recordings were performed at room temperature using Multiclamp 700B amplifier (Molecular Devices) from borosilicate patch electrodes (P-97 puller and P-1000 puller; Sutter Instruments) containing a potassium-based intracellular solution (in mM): 123 K-gluconate, 10 KCl, 10 HEPES; 1 EGTA, 1 MgCl2, 0.1 CaCl2, 1 Mg-ATP, and 0.2 Na4GTP (pH 7.2). 0.06% sulpharhodamine dye was added to the intracellular solution to confirm the selection of multipolar neurons. The extracellular solution consisted of (in mM): 140 NaCl, 5 KCl, 1.25 NaH_2_PO_4_, 1 MgCl_2_, 2 CaCl_2_, 10 glucose, and 10 HEPES (pH 7.4). Data were digitized at 10 – 20 kHz and low-pass filtered at 1 - 2 kHz. Recordings were omitted if access resistance was >30 MΩ. Whole-cell recordings were clamped at -70 mV and corrected for a calculated - 10mV junction potential. Rheobase was determined by a step protocol with 5 pA increments, where the injected current had a 25 ms duration. Action potential waveform parameters were all analyzed in reference to the threshold. Repetitive firing step protocols ranged from -20 pA to +50 pA with 5 pA increments for a duration for the isogenic KO line. This was adapted for the patient-derived iNeurons as it took more current to elicit their rheobase. The repetitive firing step protocol ranged from -40 pA to +90 pA with 10 pA increments. No more than two iNeurons per coverslip were used to reduce the variability. Data were analyzed using the Clampfit software (Molecular Devices), while phase-plane plots were generated in the OriginPro software (Origin Lab).

### Proteomic Profiling by Tandem-Mass-Tag-based Mass Spectrometry

Approximately 150 μg of total protein was extracted from 3 independent NGN2 transductions at DIV14 for the patient-derived CTRL-F1 and R607* iNeurons (total of 6 samples per patient family) using 8 M urea and 100 mM ammonium bicarbonate. These protein samples were reduced with 10 mM tris(2-carboxyethyl)phosphine for 45 min at 37 °C, alkylated with 20 mM iodoacetamide for 45 min at room temperature, and digested by trypsin (Promega) (1:50 (w/w) enzyme-to-protein ratio) overnight at 37 °C. The resulting peptides were desalted with the 10 mg SOLA C18 Plates (Thermo Scientific), dried, labeled with 16-plex tandem mass tag reagents (Thermo Scientific) in 100 mM triethylammonium bicarbonate, and quenched with 5% hydroxylamine before being pooled together. 40 μg of the pooled sample was separated into 36 fractions by high-pH reverse-phase liquid chromatography (RPLC) using a homemade C18 column (200 μm × 30 cm bed volume, Waters BEH 130 5 μm resin) running a 70 min gradient from 11 to 32% acetonitrile− 20 mM ammonium formate (pH 10) at a flow rate of 5 μL/min. Each fraction was then loaded onto a homemade trap column (200 μm × 5 cm bed volume) packed with POROS 10R2 10 μm resin (Applied Biosystems), followed by a homemade analytical column (50 μm × 50 cm bed volume) packed with Reprosil-Pur 120 C18-AQ 5 μm particles (Dr. Maisch) with an integrated Picofrit nanospray emitter (New Objective). LC-MS experiments were performed on a Thermo Fisher Ultimate 3000 RSLCNano UPLC system that ran a 3 h gradient (11− 38% acetonitrile−0.1% formic acid) at 70 nL/min coupled to a Thermo QExactive HF quadrupole-Orbitrap mass spectrom-eter. A parent ion scan was performed using a resolving power of 120 000; then, up to 30 of the most intense peaks were selected for MS/MS (minimum ion counts of 1000 for activation) using higher energy collision-induced dissociation (HCD) fragmentation. Dynamic exclusion was activated such that MS/MS of the same m/z (within a range of 10 ppm; exclusion list size = 500) detected twice within 5 s was excluded from the analysis for 30 s.

### Proteomic Data Analysis

LC-MS data were searched against a UniProt human protein database (ver 2017-06, 25020 entries) for protein identification and quantification by Protein Discover software (Thermo). Only proteins with 2 or more unique peptides were used for downstream analysis. Protein abundance quantification was normalized by taking the sum of each TMT channel and normalizing it to a control sample. Protein quantities were log2 transformed to calculate fold changes and p-values were calculated using Student’s t-test. Post UniProt accessions were converted to official Gene Symbols using UniProt’s Retrieve/ID mapping tool. The resulting data were subjected to the Benjamini-Hochberg procedure for correcting multiple hypothesis testing. We selected arbitrary cutoffs of adjusted P < 0.05 and a log2FC ± 20% for “differentially expressed proteins”. The differentially expressed protein dataset was used for principal component analysis, and pathway enrichment analysis.

### Pathway Enrichment Analysis

To gain further insight into the molecular perturbations, we used g:Profiler for over-representation analysis using the default parameters except for excluding electronic GO annotations with our differentially expressed proteins dataset (Reimand et al., 2016, 2019). We used GO terms from Biological Process categories with FDR < 0.05 from this analysis. The following data tables containing enriched GO terms were used to generate an enrichment map using the Cytoscape plugin Enrichment Map with an FDR cutoff of 0.05 (Merico et al., 2010; Shannon et al., 2003). The resulting network was first automatically annotated using the AutoAnnotate plugin to aid in the discovery of biological themes of interest (Kucera et al., 2016). The labels for closely clustered GO terms were edited for readability and comprehension using Adobe Illustrator. Clusters of interest were further examined by selecting GO terms belonging to the respective cluster and plotting the most differentially expressed proteins.

### Measurement of mitochondrial respiration

iNeurons were replated on DIV 4 onto polyornithine/laminin coated in the Seahorse XF96 Cell Culture Microplate (Agilent Technologies) at a density of 50 000/well, with the four corner wells left empty for the Seahorse XFe96 analyzer calibration. On DIV 5, primary mouse astrocytes were added at a density of 25 000/well to support iNeurons’ viability and maturation. Half media changes were performed every other day and wells were maintained until DIV 14 for recordings. At DIV 9, iNI was supplemented with 2.5% FBS which was adapted from (Zhang et al., 2013). On DIV 13, the XF extracellular flux sensory cartridge was hydrated with Ultrapure water and incubated overnight at 37°C in a CO_2_ free incubator. On DIV 14, mitochondrial stress test media was made (XF base media, 0.5 mM sodium pyruvate, 1.35 mM GlutaMAX, and 8.75mM filtered glucose) and warmed to 37°C. iNeurons were then washed twice with 200 uL of the MST media, after the second wash 180 uL of the MST media was added to the wells and left to incubate at 37°C for 1 hr in a CO_2_ free incubator. The XF extracellular flux sensory cartridge was removed and the MST assay compounds were added to final concentrations of (3 µM Oligomycin, 1 µM FCCP, and 1 µM Rotenone/Antimycin A). The XF extracellular flux sensory cartridge was then put into the Seahorse XFe96 analyzer to calibrate compounds and sensors. After, the XF96 Cell Culture Microplate was then inserted into the XF extracellular flux sensory cartridge and both plates were taken up into the Seahorse XFe96 analyzer. Measurements for each respiration phase were taken over 21-minute windows with measurements every 7 minutes, drugs were injected one after another with rotenone and antimycin A injected together. A CyQuant cell proliferation assay was performed after recordings to normalize oxygen consumption rate. Analysis was performed in the Wave software (Agilent Technologies).

### Statistical analysis

Data are expressed as mean ± SEM. Three viral NGN2 transductions were used as biological replicates for statistical analysis, except for CTRL-F2 and G1744* MEA data, where two biological replicates were used. We used the Student’s unpaired t-test, two-way ANOVA, and post hoc Sidak tests in GraphPad Prism 8 statistical software for statistical analyses. Sidak was used to correct for multiple comparisons. Grubbs’ test was used to remove outliers. The p-values in the figure legends are from the specified tests, and p < 0.05 was considered statistically significant.

